# Advances in multi-trait genomic prediction approaches: Classification, comparative analysis, and perspectives

**DOI:** 10.1101/2025.03.05.641610

**Authors:** Alain J. Mbebi, Facundo Mercado, David Hobby, Hao Tong, Zoran Nikoloski

**Affiliations:** Bioinformatics Department, Institute of Biochemistry and Biology, University of Potsdam, Karl-Liebknecht-Str. 24-25, 14476 Potsdam-Golm, Brandenburg, Germany; Systems Biology and Mathematical Modeling Group, Max Planck Institute of Molecular Plant Physiology, Am Mühlenberg 1, 14476 Potsdam-Golm, Brandenburg, Germany

**Keywords:** Genomic prediction, multi-trait, machine learning, deep learning, crop improvement, breeding

## Abstract

Traits in any organism are not independent, but show considerable integration, observed in a form of couplings and trade-offs. Therefore, improvement in one trait may affect other traits, often in undesired direction. To account for this problem, crop breeding increasingly relies on multi-trait genomic prediction (MT-GP) approaches that leverage the availability of genetic markers from different populations along with advances in high-throughput precision phenotyping. While significant progress has been made to jointly model multiple traits using a variety of statistical and machine learning approaches, there is no systematic comparison of advantages and shortcomings of the existing classes of MT-GP models. Here, we fill this knowledge gap by first classifying the existing MT-GP models and briefly summarizing their general principles, modeling assumptions, and potential limitations. We then perform an extensive comparative analysis with ten traits measured in an *Oryza sativa* diversity panel using cross-validation scenarios relevant in breeding practice. Finally, we discuss directions that can enable the building of next generation MT-GP models in addressing pressing challenges in crop breeding.

## Introduction

The food system is facing a challenge due to interrelated effects of climate change, growing world’s population, and increasing scarcity of resources [1]. Breeding of crops resilient to changes in environmental cues with little to no penalty to yield provides one way to address this challenge [2, 3]. However, resolving this problem requires understanding of the genetic and molecular mechanisms underpinning trait integration [4].

Traditional breeding requires significant resources to develop varieties with improved traits of interest [5, 6]. Moreover, breeding techniques based on marker assisted selection are less suitable for complex traits controlled by multiple quantitative trait loci (QTL) [7, 8, 9]. To mitigate these shortcomings, Meuwissen et al. proposed genomic selection (GS) [10] that leverages the advances in genotyping technologies with genomic prediction (GP) model that facilitates shortening of the breeding cycle [11, 12].

The basic principle of GS involves designing a training set (TRN) of genotypes for which both genotypic and phenotypic data are available (Figure 1). Models are then trained to predict the observed traits based on genomic data (*i*.*e*., genetic markers) using diverse machine learning (ML) approaches.

**Fig. 1.**
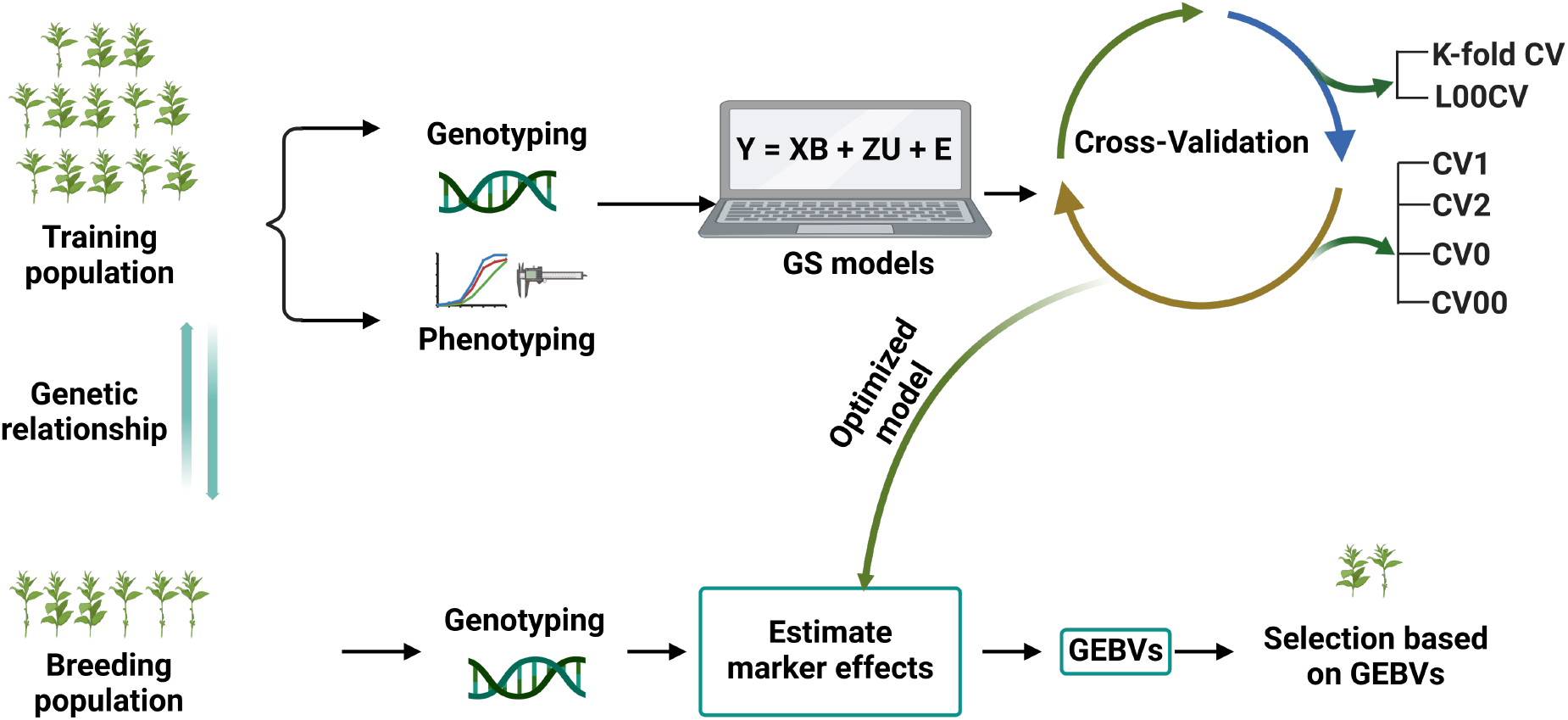
Schematic overview of genomic selection. Showcased are the main steps involved in the genomic selection (GS), starting with the collection of phenotypic and genotypic data from a training population (*e*.*g*., inbreeds or hybrids). Depending on the prediction objective and the sample size, different cross-validation (CV) schemes along with collected data are used to train the predictive models; these models are subsequently used to determine genomic estimated breeding values (GEBVs). The GEBVs are then applied to a testing population that is only phenotyped and from which individuals with desired performances are selected without the need for direct phenotyping. Briefly, in *k*-fold CV, the population under consideration is partitioned into *k* folds of approximately equal size; the model is trained on (*k −* 1) folds while the remaining fold is used for validation until each fold has been used as a validation set. LOOCV is similar to the former except for the fact that a single individual is used for validation. On the other hand, CV0, CV00, CV1 and CV2 are employed under multiple environments settings and they correspond respectively to the prediction of seen genotypes in unseen environments, unseen genotypes in unseen environments, unseen genotypes in seen environments and genotypes seen in some environments to be predicted in other seen environments. Also shown is a multivariate linear mixed effects model from which most MT-GP can be derived. In this formulation, **Y** represents the observed matrix of phenotypic values, **X** a design matrix of covariates and corresponds to the coefficient matrix of fixed effects **B**; the design matrix **Z** contains the random effects **U** and **E** is the matrix of residuals.

The resulting GP model is in turn used to predict genomic estimated breeding values (GEBVs) for a testing set or selection population (TST), for which only genotypic data are available [13]. The performance of the GP model (*i*.*e*., its prediction accuracy) computed as a correlation between GEBVs and the measured phenotypes of a trait using different cross-validation (CV) schemes. Since different ML approaches can account for both large and small QTL effects [14], GP has been successfully applied to improve trait selection across several major crop species, including: rice, wheat, sorghum, and corn [15, 16, 17, 18, 19].

Advances in high-throughput phenotyping (HTP) technologies have facilitated high precision, simultaneous measurements of multiple traits (MT) [20, 21]. As a result, these technologies have propelled the development of GP models that can predict multiple traits simultaneously. The existing single trait GP (ST-GP) models, including: ridge regression best linear unbiased prediction (rrBLUP) [22] and the so-called Bayesian alphabet [23, 24], are not appropriate to address this problem as they neglect the genetic correlations between MTs [25]. Their applications with MTs usually involve transformations of the phenotype matrix (*e*.*g*., weighted linear combination of several traits [26] and vectorization) or training of multiple STGP models, for each ST separately.

Since there is a considerable integration of traits in any organism, observed in a form of couplings or trade-offs [4], recent developments have aimed to develop GP models for multiple traits, termed MT-GP. The existing MT-GP models differ in the number of trait they can efficiently model and how population structure and marker effects are handled.

While significant progress has been made to jointly model MTs with a variety of statistical and machine learning approaches [27], there is no systematic comparison of advantages and shortcomings of the existing classes of MT-GP models. Here, we aim to fill this knowledge gap by first briefly summarizing the general principles, modeling assumptions, and potential limitations of the existing MT-GP models, followed by an outline of existing strategies used to assess the performance of MT-GP models. We then perform an extensive comparative analysis with different realistic cross-validation scenarios, using ten traits of which five are yield-related and five metabolic traits measured in an *Oryza sativa* diversity panel composed of 506 accessions characterized with genomic data on ∼900k single nucleotide polymorphisms (SNPs). Finally, we discuss possible directions that can enable the building of next generation MT-GP models in addressing pressing challenges in breeding.

## Statistical learning for MT-GP

### Main principles of MT-GP models

MT-GP models have gained significant traction over the past decade as the breeding community has sought to exploit the genetic correlation between traits to improve the accuracy and selection when simultaneously analyzing MTs [28, 29]. Unlike ST-GP models [25], that only account for between-individuals relationship and model single traits, MT-GP models use several traits simultaneously either by considering their matrix representation or a link function that jointly quantify the traits and/or their covariance. As a result, MT-GP models can boost the predictability of traits with low heritability if they are correlated with others with high heritability [30, 31, 32]. Like ST-GP, the aim of MT-GP models is to establish a mathematical relationship between MTs measured in a population of genotypes and the corresponding genomic data. When designing a MT-GP model, often two main scenarios are considered: (i) multiple groups of samples, each with a separate set of genotypes on the same set of markers and (ii) a single group of samples associated with one set of genotypes, termed multitask and multiple output learning, respectively [33]. In doing so, a single model accounting for all traits of interest is trained and the between-traits and/or between-individuals correlation are explicitly modeled.

### Assumptions of MT-GP models

The existing MT-GP models differ based on the assumptions regarding the underlying distribution of the data and the trade-offs between the resulting accuracy and interpretability.

As a result, MT-GP models can be categorized into three groups (see Table 1), namely: parametric, nonparametric, and semiparametric [68, 69]. Most MT-GP models represent a special case of a multivariate linear mixed effects model (MLMM). Therefore, we follow the notations in [34] and consider that MLMM for *s* traits recorded on *n* individuals can be written in matrix form as:

**Table 1.** Classification of multi-traits genomic prediction models. Shown is an non exhaustive list of recently developed MT-GP models with their corresponding references. The abbreviations used stand for: Stochastic search variable selection (SSVS), Multiple-trait (MT), Multiple-environment (ME), ridge regression (RR), best linear unbiased prediction (BLUP), weighted genomic BLUP (wGBLUP), Least absolute shrinkage and selection operator (LASSO), Multi-task learning (MTL), Elastic net (EN), Structured regularization with underlying sparsity (SPRING), Kernelized multivariate (KM), Partial least square (PLS), Random forest (RF), Multi-target ensemble regression chains (MTERC), Convolutional neural networks (CNNs), Recurrent neural networks (RNNs), Multi-layer neural network (MLNN), Deep learning (DL), Multi-trait Poisson deep neural network (MPDN), Reproducing kernel Hilbert space (RKHS), Support vector regression (SVR), Quasi multitask SVR (QMTSVR), Singular value decomposition (SVD), Multi-output regression stacking (MORS), Multi-target regressor stacking (MTRS), Multivariate regression with covariance estimation (MRCE), Multiple output regression (MOR) and Mega-scale linear mixed models (MegaLMM).

| Categories | Types | Models | References |
| --- | --- | --- | --- |
| Parametric | Bayesian models | Bayesian MTME | [34] |
|  |  | MT-BayesA | [31] |
|  |  | MT-BayesB | [35], [31] |
| | | MT-BayesCπ | [36] |
|  |  | MT-Bayesian rr-BLUP | [37] |
|  |  | MT-BayesSSVS | [38] |
|  |  | MT-BayesAS | [39] |
|  |  | Bayesian MORS | [40], [41] |
|  |  | Bayesian MTRS | [42] |
|  | Penalized multivariate regression models | MT-Bayesian LASSO | [35] |
|  |  | MT-LASSO | [43], [44] |
|  |  | MT-RR | [44] |
|  |  | MRCE | [45] |
|  |  | MOR | [33] |
|  |  | L<sub>21</sub>-Joint | [46] |
|  |  | L<sub>1</sub>-MTL | [47] |
|  |  | Cluster-based MTL | [48] |
|  |  | L<sub>21</sub>-MTL | [47] |
|  |  | Trace-norm Regularized MTL | [49] |
|  |  | MTL group-EN | [50] |
|  |  | MTL group-RR | [50] |
|  |  | SPRING | [51] |
|  | Mixed linear models | MT-BLUP | [52], [53] |
|  |  | MT-GBLUP | [36] |
|  |  | MT-rrBLUP | [38], [39] |
|  |  | MT-wGBLUP | [54] |
|  |  | Multivariate pedigree-BLUP and -GBLUP | [31] |
|  | Other multivariate regression models | SVD-BMTME | [55] |
|  |  | MT-PLS | [56] |
|  |  | KMLASSO | [57] |
|  |  | MegaLMM | [58] |
| Nonparametric |  | MTERC | [59] |
|  |  | CNNs | [60] |
|  |  | RNNs | [61] |
|  |  | MLNN | [62], [34] |
|  |  | MT-DL | [61], [63], [64] |
|  |  | MPDN | [65] |
| SemiParametric |  | MT-RF | [66] |
|  |  | MT-RKHS | [67] |
|  |  | QMTSVR | [67] |

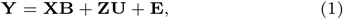

with **Y** ∈ ℝ ^*n*×*s*^, an observed matrix of phenotypic values for *s* traits on *n* individuals, that serve as responses, **X** ∈ ℝ ^*n*×*p*^ a design matrix of covariates (*i*.*e*., fixed effects) including a row of 1*s* as intercepts for each response and corresponding to the coefficient matrix of fixed effects **B** ∈ ℝ ^*p*×*s*^; clearly, each column of **B** represents the fixed effect coefficients of all covariates for a particular trait. The design matrix **Z** ∈ ℝ ^*n*×*r*^ contains the random effects **U** ∈ ℝ ^*r*×*s*^ that quantify the deviation from fixed effects such that each row of **U** represents the random effect for a single genotype for all *s* traits. Finally, **E** ∈ ℝ ^*n*×*s*^ is the matrix of residuals for the *s* traits representing the part of **Y** not explained by the model. It is further assumed that **E** ∼ *MV N* (0, **I**_*n*_, **R**) and **U** ∼ *MV N* (0, **G**, Σ_*g*_), with **G**, the genomic relationship matrix (GRM) that can be derived using the method suggested in [70]. Specifically, 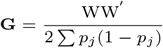, where W is the centered SNP matrix with entries *w*_*ij*_ = *m*_*ij*_ − 2*p*_*j*_, *p*_*j*_ is the allele frequency for the *j*^th^ SNP, *m*_*ij*_ the genotype coded 0, 1, and 2, denoting reference homozygote, heterozygote, and alternate homozygote genotype, for the *i*^th^ sample and *j*^th^ SNP. Further, **R** denotes the residual covariance matrix and **I**_*n*_ the *n*-dimensional identity matrix. The notation **Y** ∼ *MV N* (**M, U, V**), is used to denote that the matrix **Y** ∈ ℝ ^*n*×*s*^ follows a matrix-variate normal distribution [71] with location parameter **M** ∈ ℝ ^*n*×*s*^ (*i*.*e*., the expected value of **Y**) and covariance matrices **U** ∈ ℝ ^*n*×*n*^ and **V** ∈ ℝ ^*s*×*s*^. Deriving closed form and unique solutions analytically for the maximum likelihood estimates of covariance matrices under matrix-variate normal distribution is a challenging task, particularly for unstructured covariance and high dimensional data. In practice, this obstacle can be tackled by assuming separability of **Σ** the covariance between variables and samples, which amounts to writing **Σ** as a Kronecker product of the between and within variable covariances (*i*.*e*. **Σ** = **U** ⊗ **V**), and by imposing identifiability constraints such as tr(**V**) = *s* (*i*.*e*., the between samples covariance is assumed to be the identity matrix) [72].

### Parametric models

These type of MT-GP models rely on specific distributional assumptions with a predefined link function between traits and markers. One of the most common parametric methods is the multi-trait BLUP (MT-BLUP) [28, 52, 53], which is an extension of its ST version. To accommodate the presence of several traits and address selection bias, if correlated traits are analyzed individually [73], MT-BLUP incorporates genetic and residual correlations between traits [34]. MT-BLUP can be derived from (Eq.1) by stacking columns of the respective matrices on top of each other (*i*.*e*., vectorization) and assuming a given probability distribution for all random effects (see, for example, [38]). Under the assumption of same covariance for all loci and independent SNP effects, that is ultimately equivalent to assuming a common GRM for all traits, its genomic variant (MT-GBLUP) can be derived in a similar way [36]. By extending the ST-rrBLUP constant variance assumption to constant covariance for all SNPs, its MT version (*i*.*e*., MT-rrBLUP) [39] was also proposed. In a Bayesian framework, covariance between SNPs across traits was explicitly accounted for through the introduction of latent variable in the expectation of random effects. Because of the Bayesian Markov chain Monte Carlo(MCMC) approach used for parameters estimation, this model is also referred to as MT-Bayesian rrBLUP.

This rather strong assumption of common G for all traits was further relaxed in [54] to obtain a weighted multi-trait GBLUP (MT-wGBLUP) model in which a sparse latent variable model approach is used to estimate breeding values and SNP effects under heterogeneous SNP-covariance between genomic regions (*e*.*g*., chromosomes or group of SNPs) [74].

Nonetheless, the number of parameter to be estimated can quickly increase leading to a computational burden and unreliable estimates for unstructured covariance. As a remedy, MT-Bayesian models have been proposed that utilize prior distributions to incorporate prior knowledge on marker effects and handle different types of genetic architectures more flexibly [75]. We refer the interested reader to [34] for detailed explanations on how the Bayesian variants of the MT-BLUP and related models can be derived with proper priors assigned to the parameters of interest.

Additionally, to account for departure from normality, often exhibited by some traits, MT extensions of the Bayesian alphabet including: BayesA, BayesB, BayesC*π*, Bayesian LASSO and Bayesian rr-BLUP have also been proposed (see, [31, 35, 37]). For instance, MT-BayesC*π* estimates the marker effects by variable selection and assumes that each locus can have an effect on any combination of traits [36].

Another model called MT-BayesAS assumes similar genomic covariance structure for SNPs within a given genomic regions, but different genomic covariance for SNPs in different genomic regions, and relies on multi-trait random regression model to explicitly model heterogeneous variance and covariance as a latent variables model [39, 74]. Additionally, MT-BayesA and MT-Bayesian LASSO that incorporate the assumption of heterogeneous covariances could also be derived from MT-BayesAS.

When information about multiple environments is available as it is often the case for genetic evaluation, the above MT models cannot account for genotype-environment (G x E) and trait-genotype-environment (T x G x E) interactions. The Bayesian whole genome prediction [29] that makes use of an efficient MCMC procedure for parameter estimation with an exact Gibbs sampling for the posterior distribution was pioneered to fill this knowledge gap [76, 25]. For instance, several traits for maize and wheat have been predicted in diverse environments using Bayesian multiple-trait multiple-environment (BMTME) models [55]. This set of approaches incorporates G x T and T x G x E interactions to improve predictabilty. A variant of BMTME that is equivalent to principal component analysis and termed SVD-BMTME, handles correlated traits by making use of singular value decomposition (SVD) on the trait matrix. This is performed to remove the information related to other covariates (*i*.*e*., de-correlation). Since the uncorrelated and decomposed vectors are subsequently used as traits, the model implementation becomes straightforward as computational tools designed for univariate models can be employed. The final MT predictions are derived by transforming back the decomposed vectors to their original scale (*i*.*e*., before decomposition) [55].

In a different perspective, models accounting for complex covariance structures in simultaneous analysis of MT in several environments have also been proposed using a two-stage approach. The first stage uses the same ST-GP model to predict each trait. The predicted traits are then used in the second step as predictors in a MT model for final predictions. These are built upon a Bayesian extension of multi-target regressor stacking initially proposed in [42] and include variants of the Bayesian multi-output regression stacking (BMORS) [40, 41]. This framework present flexibility in term of modeling depending on the data at hands, whereby the user can account for both linear and non-linear MT relationships in the second stage by selecting the desired model in the first stage. In this regards, random forest regression has been employ in the first step [27] to account for non-linearity.

These MT models are implemented as their ST versions, whereby vectorizations of the matrix of traits is applied; in the Bayesian setting, these models can quickly become computationally expensive because of the MCMC steps required during parameter estimation. To address this issue, multivariate regression models that incorporate different correlation structures to better exploit the possible shared between-traits and between-genotypes relationships have been proposed [45, 33, 46]. These models are built upon (Eq. 1) and do not explicitly model genetic random effects (*i*.*e*., **Y** = **XB** + **E**). Instead, with suitable distributional assumption on the residuals, they use a multivariate likelihood framework and impose different regularization on parameters of interest (*i*.*e*., regression coefficients and covariance matrix). Under the assumption that the covariance is the identity matrix, these models can be further simplified and design as simple regularized multiple output linear regression. These include the multivariate extension of least absolute shrinkage and selection operator (LASSO), ridge regression and elastic net [44]. To account for possible non-linear dependence between the predictors and responses, Kernelized multivariate LASSO [57] that solves 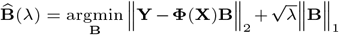, with kernel function Φ, has also been proposed and applied.

Another challenge often encountered in data used for GP is related to the derivation of parameter estimates unaltered by variables collinearity. In a multivariate regression framework, this has been explored using the MT equivalent of partial least square regression [77] (MT-PLS) and applied to enhance prediction accuracy of potato cultivars in new environments [56]. MT-PLS has the ability to simultaneously accommodate different factors in small-*n* large-*p* setting and with MTs. Without the random effect part, MT-PLS can be derived from (Eq.1), the only difference with the previously mentioned multivariate regression being that **Y** is regressed on latent variable or scores **T** instead of the original predictors **X**. The estimate is derived in an iterative procedure that maximizes the covariance between the original traits and the latent variables, and the final coefficients rotated or converted back to the original space of **X**.

### Nonparametric models

Unlike their parametric counterparts, nonparametric approaches do not rely on strict distributional assumptions, rendering them more robust to outliers or when the normality assumption is violated, as it is often the case with experimental data sets. Further, in nonparametric approaches, there is no predefined link function between phenotypes and genetic markers. Instead, this relationship is learned from the data. This class of models has recently gained popularity because of their flexibility and ability to reduce modeling bias [78, 69].

Recently developed approaches include the multi-target ensemble regression chains that selects assistant trait automatically and predict the genomic estimated breeding values of the target trait using genotypic information only [59]. MT deep learning (MT-DL) approaches, including: (i) convolutional neural networks (CNNs), that can capture spatial dependencies between markers in modeling effects on phenotypic traits, provide hierarchical feature representations, and leverage large-scale genomic data; (ii) recurrent neural networks (RNNs), useful for modeling of time-series data; and multi-layer neural network (*i*.*e*., input, hidden and output layers) [62] (MLNN) that bypasses the restrictive assumption of additive genetic effects of markers. More precisely, the input layer consists of SNPs, the neurons or mapping units in which the weighted sum of nodes from the SNPs is computed constitute the hidden layer and the output layer contains the model outcome. Nonetheless, for large number of SNPs, the MLNN can quickly become computational intractable requiring the usage of a subset of SNPs (*e*.*g*., significant SNPs derived from MT-genome wide association studies). In particular, they have shown great promise thanks to their ability to model highly non-linear relationships and interactions among genetic markers [60].

Extension to this work aimed to account for MT interactions in multiple environments [61] and different type of responses (*e*.*g*., binary and ordinal) [79] have also been studied. Using three real data sets comprising elite maize and wheat lines, it was demonstrated that MT-DL model is less computational expensive and is a competitive alternative to the BMTME, with the best predictions achieved when the G x E interaction was ignored. In contrast, the BMTME model outperformed the MT-DL in terms of prediction accuracy when G x E interaction was considered.

The non-parametric nature of deep learning and related approaches allows the model to learn without predefined assumptions about the relationships between traits, making it ideal for handling complex trait architectures. Their flexibility can facilitate the incorporation of multi-environment and multi-trait data simultaneously, providing robust predictions across different conditions. Nonetheless, the application of MT-DL models in genomic prediction is still facing some challenges. They are more difficult to generalize, as they require larger sample size compared to the parametric counterparts [64], they are unable to estimate the between-traits covariance, and are often computationally expensive in terms of parameter tuning.

### Semi-parametric models

Blending parametric and non-parametric machinery, semi-parametric approaches have also emerged as a powerful tool for MT-GP. Previously used in ST-GP [80] and relying on kernel functions to build a covariance structure between traits, kernel-based regressions such as support vector (SVR) and reproducing kernel Hilbert space (RKHS) have also been adapted to accommodate MT-GP. These adaptations include: (i) the quasi multitask SVR (QMTSVR) with hyperparameter tuning derived from genetic algorithm, (ii) its weighted alternative and (iii) the MT-RKHS, where the usual G matrix is replaced with a nonlinear kernel K from SVR [67]. RKHS method is particularly interesting as it leverages kernel functions to capture complex, non-linear relationships between genetic markers and traits while still maintaining some parametric structure. This method has been shown to provide more accurate predictions than purely parametric models, especially when dealing with complex traits that exhibit non-linear genetic effects.

## Comparative analysis with selected MT models

### Models used in the comparative analysis and performance assessment

In what follows, the performance of the baseline ST genomic best linear unbiased prediction (ST-GBLUP) [70], the genomic variant of the rrBLUP is contrasted with that of five representative of previously discussed MT models including: MT Bayesian multi-output regressor stacking (MT-BMORS) [25], MT Multi output regression (MT-MOR) [33], MT Singular value decomposition (MT-SVD) [55], MT partial least square regression (MT-PLS) [56] and MT deep learning (MT-DL) [81]. It is worth noting that the selection of these representative MT models was based on the availability of implementation tools in the R programming language and to avoid as much as possible inclusion of models that have been assessed together in a previous comparative analysis with ST-GP. We note that our analysis differs from those presented in other studies [82, 83, 37, 43] based on the number of considered MT models across different types and the number of traits investigated.

Performances of the considered five MT-GP approaches along with that of the baseline ST-GBLUP were quantified by the Pearson correlation coefficient between predicted and observed trait values in the testing set. Final prediction accuracy values were then computed as the average performances over 100 (*i*.*e*., 20 repetitions of 5−fold CV) and 200 (*i*.*e*., 20 repetitions of 10−fold CV) for three cross-validation scenarios: (i) in CV-A, models were trained on Indica and Japonica to predict respectively Indica and Japonica. (ii) CV-B corresponds to the CV scenario where the contending models were trained on data from Indica and were used to assess the performance on Japonica and vise versa. (iii) Finally, CV-C is concerned with a random splitting with varying proportion of combination of Indica/Japonica samples to predict the remaining mixed samples of Indica and japonica accessions. Note however that, for MT-DL a controlled random CV was used, whereby instead of using a purely random split, we considered as testing set one fold from the previous 5− or 10−fold CV and the remaining folds as training set, and implemented following an adaption of the R script provided in [81].

### Phenotypic and genotypic data

To contrast the predictability of selected approaches, we used freely available phenotypic and genotypic data from previous studies [84, 85]. The number of traits that can be accommodated by a MT-GP depends on the type of model under consideration. While some [58] can effectively model thousands of traits, the computational complexity of others becomes a challenge (*e*.*g*., estimation of large covariance matrix) as the number of traits increases [81]. These challenges call for adequate planning when choosing the number of traits for a given breeding objective, to ensure optimal performance. Therefore, a fair comparative analysis dictates using the number of traits (*i*.*e*., ten in our case) that can be handled by all models.

#### Genomic data

A diversity panel of 533 *Oryza sativa* accessions, including both landraces and elite varieties, was obtained from various sources [86, 87, 88]. We obtained the bed format, together with the corresponding bim and fam files associated with the rice accessions in RiceVarMap2 database [89]. We then built a preprocessing pipeline in PLINK [90, 91] to select genotypes and variants of interest. Loci were filtered by utilizing a moving window of 100 gbp with a step of 10 gp while considering a quality threshold of 0.95. Selecting accessions for which traits information were available and removing SNPs with minor allele frequency (MAF) less than 5%, yielded a final dataset of 506 accessions and 973275 markers (see Supplementary File 1).

#### Yield-related traits

Field trials for yield-related traits were conducted in three environments: Huazhong Agricultural University, Wuhan, China (*i*.*e*., 2011 and 2012), and Lingshui County, Hainan Island (*i*.*e*., 2011). Rice seedlings were transplanted to fields in a randomized complete block design with two replications, and yield was measured from five plants per plot. This trait dataset contains rice genotypes from different populations (and subpopulations), including: Aromatic, Aus, Indica and Japonica. Detailed information on the population structure at the genotype level is provided in (Supplementary Table 1). The selected yield-related traits [92] include: yield, plant (PH) height, grain weight (GW), heading date (HD), and panicle seed setting rate (PSSR). Accession-specific values for each trait are provided in (Supplementary Table 2) whereas the min-max scaled traits are available in (Supplementary Table 3).

#### Metabolic traits

The metabolomic data set [84] includes 840 metabolites and replicates measured across 506 accessions as shown in (Supplementary Table 4). Metabolic traits in the assembled dataset come from a variety of classes including flavonoids, terpenes, fatty acids, amino acids, nucleic acid derivatives, polyphenols and phenylamines. For the metabolite selection, marker-based heritability [93] were computed for each trait. Metabolites were then selected to represent high (*h*^2^ > 0.7) and low (*h*^2^ < 0.5) heritability (see, Supplementary Table 5) and to ensure that they are representative of subpopulations present in the selected focal traits. To this end, a tricin derivative (*i*.*e*., spectra peak labeled mr1246) and C-pentosyl-apigenin O-p-coumaroylhexoside (*i*.*e*. mr1234), a flavonoid compound weighing around 711 Daltons with molecular formula C_35_H_34_O_16_ [94] were retained as high-heritability traits (*i*.*e*., *h*^2^ = 0.89). Subsequently, a tricin derivative with a spectral peak labeled as mr1198 [94] (*h*^2^ = 0.45), a polyphenol named N-Feruloyltyramine (*h*^2^ = 0.31), a compound weighing around 314 daltons with molecular formula C_18_H_19_N_*O*4_ (*i*.*e*. mr1268) [95] and LPC(1-acyl 18:2) with a spectral peak labeled mr1418 (*h*^2^ = 0.24) a fatty acid (*i*.*e*., C_26_H_50_N_07_P) [95] as low-heritability traits.

#### Trait summary, heritability, and correlation

BLUP for metabolic traits were computed using a LME with genotypes and replicates as random effects. The above SNP data was used as input in TASSEL [96], to derive the genomic relationship matrix (GRM). The obtained GRM along with traits values were then used to derive variance components in a restricted maximum likelihood (REML) framework [97] using the heritability package [98] in R statistical software (R Core Team 2021) [99].

## Performance assessment on representative MT-GP models

The comparative analysis of prediction accuracy across representative five multi-trait (i.e. MT-BMORS, MT-MOR, MT-SVD, MT-PLS, MT-DL) and a baseline single-trait (ST-GBLUP) models highlights distinct trends across different traits and cross-validation (CV) scenario.

Our findings in (Fig. 2) clearly show the influence of CV schemes on prediction accuracy. However, no consistent trend could be observed across traits and models for CV-A, where the models are trained and tested within the same subspecies (Indica or Japonica) (Fig. 2(a)-(b)), as well as CV-B, where models trained on one subspecies (Indica or Japonica) (Fig. 2(d)-(e)) are validated on the other. However, for most traits the highest prediction accuracies are observed in CV-A. This can be attributed to the genetic homogeneity within each subspecies, which simplifies the prediction task. Conversely, in CV-B, accuracies drop substantially likely due to the genetic divergence between subspecies (*i*.*e*., training and testing populations), reflecting the expected challenges of transfer learning across genetically divergent groups in GP. CV-C, which involves random splits across combined Indica and Japonica (Fig. 2(c)) sub-populations, seems to reconcile this discrepancy, highlighting the benefit of using heterogeneous training set for predictions on mixed testing populations. These observations are in line with findings in a previous study [100], emphasizing the importance of using heterogeneous training data for robust genomic predictions.

**Fig. 2.**
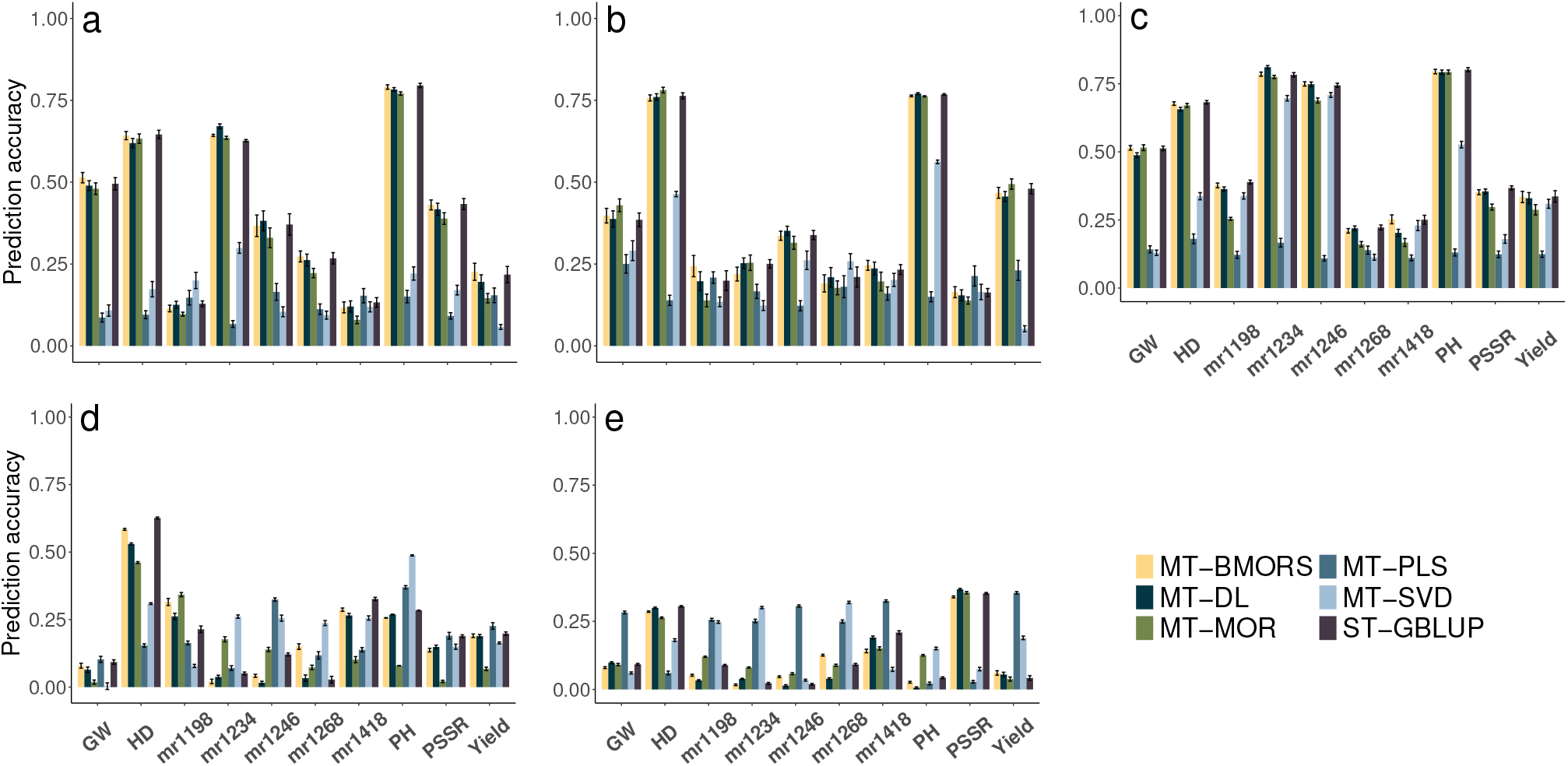
Comparison of predictabilities for MT and a baseline GP methods with a rice data set. We used 5 MT-GP models, namely: MT Bayesian multi-output regressor stacking (MT-BMORS), MT Multi output regression (MT-MOR), MT Singular value decomposition (MT-SVD), MT partial least square regression (MT-PLS) and MT deep learning (MT-DL), and single-trait genomic best linear unbiased prediction (ST-GBLUP) to predict the levels of five metabolites (*i*.*e*. mr1198, mr1234, mr1246, mr1268 and mr1418; see (Metabolic traits) section for full description) as well as five yield-related traits (*i*.*e*. yield, grain weight (GW), heading date (HD), panicle seed setting rate (PSSR) and plant height(PH)). The predictability is computed as the average Pearson correlation coefficient between observed and predicted values for the ten traits in the validation set, based on 20 repetitions of 5- and 10-fold cross-validation for respectively CV-A (panels **a** and **b**), CV-B (panels **d** and **e**) and CV-C (panel **c**). The average accuracy obtained from repeated cross-validations are reported as the height of the bars along with the standard errors. Panels **a** and **b** correspond to to the CV schemes in which models were trained on Indica and Japonica to predict traits in Indica and Japonica accessions, respectively. In contrast, panels **d** and **e** correspond respectively to the CV scenario where the models were trained on data from Indica (Japonica) and used to predict the performance on Japonica (Indica). Finally, panel **c** is concerned with the random split with varying proportion of combined Indica/Japonica samples to predict the remaining mixed samples of Indica and japonica.

Except for MT-PLS, that exhibits the lowest accuracy, MT models slightly outperform the ST-GBLUP model in most scenarios, particularly in CV-A and CV-C, where trait correlations can be effectively leveraged. Traits such as grain weight (GW) and plant height (PH) show the most significant gains with MT models, reflecting their ability to exploit shared genetic architectures. However, in CV-B, where the training and validation sets are derived from divergent genetic backgrounds, the advantage of MT models diminishes slightly, likely because of the reduced utility of trait correlations under such conditions. Similar patterns have been reported in literature [83] highlighting the utility of MT approaches in scenarios with high genetic and phenotypic correlations. Notably, MT-DL and ST-GBLUP consistently exhibit high predictive accuracy, indicating their robustness.

Between the MT models, MT-DL and MT-BMORS consistently achieve the highest prediction accuracies across all CV designs and traits. The strength of MT-DL resides in its ability to handle complex and non-linear relationships, while MT-BMORS benefits from its ability to account for trait correlations. Despite being less competitive than MT-DL and MT-BMORS for most traits, the performance of MT-MOR is not negligible in comparison to that of MT-PLS and MT-SVD. The poor performance and strong variability exhibited by MT-PLS and MT-SVD could suggest their sensitivity to specific trait architecture or a possible loss of information during the decompositions involved in the respective algorithm. Similar findings have previously been reported [81, 60] showcasing the potential of deep learning and Bayesian approaches for genomic prediction, particularly when simultaneously modeling multiple traits.

Overall, our findings underscore the value of multi-trait models in leveraging trait correlations, particularly for complex or highly polygenic traits. While single-trait models like ST-GBLUP remain effective and competitive in certain cases, MT approaches provide a significant advantage, especially when dealing with correlated traits or leveraging shared genetic architectures across traits. The consistent superiority of MT-DL and MT-MOR across traits and CV schemes suggest their potential utility in breeding applications. However, the choice of model should also consider computational challenges and trait-specific requirements, since MT-DL may demand higher computational resources compared to models like MT-BMORS or ST-GBLUP. It could be interesting for future research to investigate how MT-GP approaches could further be optimized for the benefit of practical breeding, and to also explore the integration of different data types (*e*.*g*., environmental, HTP and transcriptomic) to enhance prediction accuracy and applicability.

## Challenges and future directions

Our overview of newly developed MT-GP models and the comparative performances of parametric, semi-parametric, and non-parametric models revealed that no single approach is universally superior; rather, the choice of model often depends on the specific traits and their genetic architecture. Likewise, a similar conclusion could be reached with respect to the selected approaches used in the present analysis. Parametric models like MT-GBLUP and Bayesian methods are advantageous due to their simplicity and interpretability, particularly when the genetic architecture is well understood. Semi-parametric approaches such as MT-RKHS on the other hand, provide a balance between flexibility and interpretability, making them suitable for traits with moderate complexity. Non-parametric methods, including all variants of DL, excel in capturing intricate non-linear relationships but may require larger datasets and greater computational resources.

The future of MT-GP lies certainly in the integration of these diverse approaches, potentially through ensemble methods that combine the strengths of parametric, semi-parametric, and non-parametric models. Advances in computational power and algorithm development will further enhance the applicability and accuracy of these models. Moreover, as the availability of multi-omics data increases, integrating MT-GP with other layers of biological information, such as transcriptomics and metabolomics, promises to revolutionize our ability to predict complex traits with high precision [101, 102]. As highlighted in [61], the choice of appropriate hyperparameters for the implementation of nonparametric models such as MT-DL remains a challenge. This calls for the consideration of different network architectures and sets of hyperparameters that all together could enhance the reliability of the approach and full incorporation into practical breeding.

HTP technologies have significantly advanced the capacity to capture large-scale, multi-dimensional phenotypic data, which is critical for MT-GP. By enabling the measurement of MT across diverse environmental conditions with greater precision, HTP could enhance the accuracy of trait heritability estimates and facilitate the identification of G x E interactions. This rich data collection allows MT-GP models to better account for the complex relationships between MT and environmental factors, and could ultimately improve predictive power and selection efficiency in plant breeding programs [103]. A notable example is a study by [64] in which three traits were evaluated in 43 environments across several water regimes, showing that, despite the added complexity from the data, performance of MT deep learning model matches that of GBLUP. In addition, the ability to gather data across different growth stages and environmental conditions also supports more robust modeling of the temporal dynamics of phenotypic expression.

Nonetheless, the improved accuracy exhibited by MT models when using multi-dimensional phenotypic data from HTP should be taken with caution. While offering model flexibility the availability of these large scale datasets constitutes a challenge on its own, thereby one needs through some pre-processing, genetic correlation or heritability to select a representative subset of traits to be analyzed. Even for high genetic correlation, it has been shown that multivariate models may not be the best approach when predicting lines with small genetic relatedness [83]. Systematic assessment of MT model for different combination of heritability and genetic correlation should be keep in mind during traits selection and model evaluation. For instance, simultaneous modeling for: (i) traits with low heritability and high genetic correlation with others and (ii) traits with high heritability and high negative genetic correlation with others [104].

Another critical limitation faced by traditional MT-GP models is the number of traits they can accommodate while preserving their natural multidimensional structure and their possible shared relationships. These MT models have difficulties to cope with increasing trait numbers due to computational demands (*e*.*g*., MT-DL) and poor estimate of high-dimensional covariance matrix (*e*.*g*., regularized multivariate regression in the maximum likelihood framework). Additionally, many MT-GP models rely on complete phenotypic datasets, reducing their effectivity in partially observed traits setting. Mega-scale linear mixed models (MegaLMM) [58] that has been shown to handle thousands of traits simultaneously overcomes this constraint through the usage of sparsity-inducing priors while maintaining computational efficiency and improved prediction accuracy.

One of the key challenges in MT-GP, as for their ST counterpart, is the transferability of predictive models across environments, thereby models trained in one environment often struggle to predict trait performance accurately in a different environment due to differences in environmental conditions, G x E interactions, and trait plasticity. While MT models can leverage shared genetic architecture among correlated traits, they still face difficulties when the environmental context changes, as the model might capture trait associations that are specific to the training environment. This limitation, akin to what is observed in ST-GP [105], presents a significant challenge for breeding programs targeting broad environmental adaptability. MT multi-environment [25] and transfer learning [106, 107] GP models have been proposed to address this issue, nonetheless their applications in GP remains underdeveloped.

While time series methods have shown potential for phenomic prediction by capturing temporal patterns in trait development [63], their application in MT-GP remains largely unexplored. Incorporating time-series data could provide a more comprehensive understanding of how MTs co-evolve over time and under different environmental conditions, thereby improving the predictive accuracy and offering new insights into complex trait inter-dependencies. This approach could be particularly valuable for many agronomically relevant traits, including growth and yield potential, that are dynamic throughout the plant life cycle.

The integration of genome wide association studies (GWAS) into GP models provides another means to improve the biological interpretability of genomic prediction models [108]. GWAS can identify genetic variants associated with multiple traits, allowing for the inclusion of trait-specific marker effects in prediction models. By leveraging GWAS results, GP models can be refined to account for pleiotropic effects, (*i*.*e*., a single locus influences multiple traits), thereby improving the accuracy when predicting correlated traits. However, integrating GWAS data into MT-GP prediction models remains challenging due to the complexity of polygenic traits and the large number of genetic markers involved.

Despite successful applications across multiple model organisms and practical breeding programs, MT-GP models are prone to overfitting and a decrease of prediction accuracy has been observed with respect to their ST alternative in several comparative analysis. This make model selection a challenging task especially when secondary traits measured on genotypes from the testing population are used to predict focal traits [109]. Additionally, results from multi-locus shrinkage modelling revealed that the majority of agronomic traits in crops exhibit considerable polygenicity due to small effects of multiple QTL [110, 111, 112]. Choosing the appropriate MT model to accurately predict such traits while simultaneously accounting for the small-*n* large-*p* problem remains a challenge in MT-GP. This calls for appropriate and unbiased methodology such as the commonly used cross-validation (CV) [113, 114] for performance (*i*.*e*., predictability or prediction accuracy) evaluation. However, CV strategies in MT-GP must be adapted to emulate realistic plant-breeding mechanisms, account for the targeted prediction scenarios and insure independence between the training set and the testing one. These challenges are addressed by using CV1 and CV2 [105] corresponding respectively to the prediction of untested lines (newly developed genotypes) in tested environments and sparse testing where lines tested in some environments are to be predicted in other tested environments.

Finally, addressing the increasing global food demand in the context of a growing population is one of the most pressing challenges in plant breeding, thereby crop production must increase significantly to ensure food security. This challenge is compounded by the effects of climate change, which threaten to reduce agricultural productivity. One way to accelerate crop improvement that enables the selection of plants with optimal combinations of traits, such as higher yield and disease resistance is the usage of MT-GP models. However, the complexity of breeding for multiple traits simultaneously, especially under changing environmental conditions, presents significant obstacles. Advances in computational methods, along with the integration of multi-environment genomic prediction [13], will be crucial to addressing these challenges and ensuring a resilient global food supply.

## Key points

- We provided a classification of computational approaches for multi-trait genomic prediction.
- We pointed the differences of the underlying principles, assumptions, and potential limitations of the three classes of computational approaches for multi-trait genomic prediction.
- We compared the performance of five representative approaches for genomic prediction of ten traits related to yield and metabolism in a rice diversity panel.
- We observed a consistent superior performance of multi-trait deep learning as well as multi-output regression across the studied traits and cross-validation schemes.
- Nevertheless, the choice of approach to be used in practice depends on several factors, including the traits considered, the population structure, and genetic architecture of the traits.

## Author biographies

**Dr. Alain J. Mbebi** is a postdoctoral fellow in the Bioinformatics Department at the Institute of Biochemistry and Biology of the University of Potsdam and guest scientist in the Systems Biology and Mathematical Modeling Group at the Max Planck Institute of Molecular Plant Physiology. His current research interests include high-dimensional covariance estimation, mathematical statistics, gene regulatory networks and genome wide association study.

**Facundo Mercado** is a Master student in the Bioinformatics Department, Institute of Biochemistry and Biology of the University of Potsdam. His current research interests include computational and systems biology, genomic data analysis and parameter estimation.

**David Hobby** is a Ph.D. candidate with the Bioinformatics Department, Institute of Biochemistry and Biology of the University of Potsdam. His main research interests include development and application of machine learning approaches for time series analysis in genomic prediction.

**Dr. Hao Tong** is a postdoctoral fellow in the Bioinformatics Department at the Institute of Biochemistry and Biology of the University of Potsdam and guest scientist in the Systems Biology and Mathematical Modeling Group at the Max Planck Institute of Molecular Plant Physiology, with research interest in Statistical genetics modelling, quantitative genetics models for phenotype prediction and causal gene discovery.

**Prof. Dr. Zoran Nikoloski** is the Chair of the Bioinformatics Department at the Institute of Biochemistry and Biology at the University of Potsdam and Cooperative Research Group Leader in Systems Biology and Mathematical Modeling with Max Planck Institute of Molecular Plant Physiology. His main areas interests include: data-driven qualitative and quantitative modelling of genome-scale metabolic and gene-regulatory networks, analysis of evolutionary and optimisation processes in biological networks, and characterization of system’s functions emerging from molecular interactions.

## Data availability

We implemented all statistical models using R programming language and the codes can be freely accessed from https://github.com/alainmbebi/MT_Review. Phenotypic and genotypic data used in this study were from [84] and are available for query from reference within.

## Author contributions statement

A. J. M. - method implementation, method testing, data analysis, figure preparation, manuscript writing, F. M. - method implementation, data analysis, D. H. - method implementation, data analysis, H. T. - method implementation, data analysis, method testing, Z.N. - conceptualization, data analysis, funding acquisition, manuscript writing

